# Pharmacologic activation of Δ133p53α reduces cellular senescence in progeria patients-derived cells

**DOI:** 10.1101/2025.07.28.667224

**Authors:** Sebastien M. Joruiz, Delphine Lissa, Natalia Von Muhlinen, Patricia K. Dranchak, James Inglese, Izumi Horikawa, Curtis C. Harris

## Abstract

**Background:** Patients with Hutchinson-Gilford progeria syndrome (HGPS) show accelerated aging phenotypes and have shortened lifespan, with implications in physiological aging processes as well. While therapeutic approaches targeting the disease-causing abnormal protein, progerin, have been developed, further efforts to explore mechanistically distinct and complementary strategies are still critical to better treatment regimens. We previously showed that lentiviral vector-driven expression of Δ133p53α, a natural inhibitory isoform of p53, rescued HGPS patients-derived fibroblasts from early entry into cellular senescence, which is a downstream event of progerin-induced DNA damage. We also performed a quantitative high-throughput screen (qHTS) of approved drug and investigational agent libraries, leading to the identification of celastrol and AZD1981 as compounds that upregulate Δ133p53α protein levels.

**Methods:** To investigate whether celastrol and ADZ1981 upregulate endogenous Δ133p53α in HGPS-derived fibroblasts and reduce their senescence-associated phenotypes, we performed western blot assays (Δ133p53α, progerin, and p21^WAF1^, which mediates p53-induced senescence and is inhibited by Δ133p53α), senescence-associated β-galactosidase (SA-β-gal) staining, enzyme-linked immunosorbent assay (IL-6, which is a proinflammatory cytokine secreted from senescent cells), and qRT-PCR assays (p21^WAF1^ and IL-6).

**Results:** Treatment with celastrol (0.1 µM for 24 h) or AZD1981 (10 µM for 24 h) reproducibly increased Δ133p53α expression and decreased p21^WAF1^ expression in two strains of fibroblasts derived from HGPS patients. These compounds reduced the percentage of SA-β-gal-positive senescent cells and the secretion of IL-6 into culture medium in both of these fibroblast strains, irrespective of their different basal levels of senescence and IL-6 secretion. These compounds had no effect on the level of progerin.

**Conclusion:** Celastrol and ADZ1981 upregulate endogenous Δ133p53α and, reproducing the effects of its vector-driven expression, inhibit cellular senescence and IL-6 secretion in HGPS-derived fibroblasts. Their progerin-independent action suggests that they may synergize with currently available progerin-targeting therapies. This study also warrants further investigation of these compounds for potential applications in other diseases and conditions in which Δ133p53α-regulated senescence plays a role.

## INTRODUCTION

The human *TP53* gene encodes not only the full-length protein consisting of 393 amino acids (simply referred to as “p53” in general and in this manuscript as well) but also at least a dozen of naturally occurring isoforms due to alternative RNA splicing and/or alternative usage of transcription and translation initiation sites [1]. Among these *TP53*-encoded isoforms, Δ133p53α is an N-terminally truncated isoform that is generated via an alternative transcription initiation from intron 4 and an alternative translation from the methionine codon 133 [1, 2]. In various types of normal human cells, the expression level of Δ133p53α is primarily regulated via protein turnover involving chaperone-assisted selective autophagy [3-5], which leads to diminished expression of endogenous Δ133p53α upon cellular senescence [2, 3, 5-9]. The retroviral or lentiviral vector-driven expression of Δ133p53α prevents or delays normal human cells from entering cellular senescence [2, 3, 5, 6, 8, 9], supporting the direct role of Δ133p53α in regulating human cell senescence. Of particular importance, Δ133p53α-expressing cells, not entering senescence, exhibit their inherent normal functions, such as neuroprotective activity of Δ133p53α-expressing astrocytes [5, 8-10] and anti-tumor activity of Δ133p53α-expressing T lymphocytes [11]. Although inhibition of p53 activities could lead to genome instability and increased oncogenesis [12, 13], Δ133p53α preferentially inhibits p53-mediated senescence in a dominant-negative manner, while maintaining or even enhancing p53-mediated DNA repair [7-9, 11, 14, 15]. This selective inhibitory nature of Δ133p53α suggests a potential therapeutic value of Δ133p53α in senescence-associated diseases and conditions with minimum risk of genome instability or oncogenesis. In an attempt to pharmacologically activate Δ133p53α for future therapeutic applications, we performed a cell-based quantitative high-throughput screen (qHTS) on approved drugs, investigational agents and chemical probe libraries using fluorescently labeled Δ133p53α protein, which identified two compounds (celastrol and AZD1981) that upregulate Δ133p53α at the protein level in normal cells [16]. Celastrol, a pentacyclic quinone methide, is a natural product used in traditional Chinese medicine. The molecule has multiple molecular mechanisms, likely due to its thiol reactivity [17-19]. These activities include acting as an Hsp70 inducer and proteasome inhibitor [20, 21]. AZD1981 is a prostaglandin DP2 receptor antagonist developed for asthma [22].

Hutchinson-Gilford progeria syndrome (HGPS) is a genetic disease caused by a mutation in the *LMNA* gene, which produces an abnormal nuclear envelope protein called progerin [23]. Patients with HGPS exhibit premature aging phenotypes in childhood and have a short lifespan, with an average life expectancy of 13-15 years [24]. Although the occurrence of HGPS is very rare, this disease also has significant implications for natural aging processes at both the organismal and cellular levels, as well as for interventions against aging in healthy individuals [25, 26]. Similar to the premature aging phenotypes observed in patients with HGPS, their derived cells in culture (often fibroblasts) display a premature onset of cellular senescence accompanied by proinflammatory cytokine production, which is attributed to progerin-induced accumulation of DNA damage and hyperactivation of p53 [27-29]. Our previous study showed that lentiviral expression of Δ133p53α delayed the onset of cellular senescence and reduced the production of the proinflammatory cytokine IL-6 in fibroblasts derived from HGPS patients, concurrent with mitigated DNA damage and repressed expression of p21^WAF1^, a p53-inducible gene that mediates cellular senescence [7]. In this study, we investigate whether treatment with celastrol or AZD1981 can reproduce the effects of lentiviral expression of Δ133p53α in HGPS-derived fibroblasts, exploring the potential therapeutic value of these compounds in HGPS.

## METHODS

### Cells and cell culture

A fibroblast strain AG11513, from an 8-year-old female HGPS patient, was obtained from Coriell Institute for Medical Research (https://catalog.coriell.org/). Another fibroblast strain HGADFN271, from a 1-year-3-month-old male patient, was obtained from Progeria Research Foundation (https://www.progeriaresearch.org/). These cells were grown in DMEM medium (Corning #15-013-CV) supplemented with 10% fetal bovine serum (Sigma-Aldrich #F0926), 1% L-glutamine (Thermo Fisher #25030081) and 1% penicillin/streptomycin (Thermo Fisher #10378016) at 37 °C under a humidified atmosphere of 5% CO_2_. Cells were passaged at a split ratio of 1:3 (AG11513) or 1:4 (HGADFN271).

### Compounds and treatment

Celastrol was purchased from Cayman Chemical (#70950) and dissolved in DMSO as 100 µM stock solution. AZD1981 was also purchased from Cayman Chemical (#20763) and dissolved in DMSO as 10 mM stock solution. Cells were treated with celastrol at a final concentration of 0.1 µM, AZD1981 at a final concentration of 10 µM, or DMSO vehicle control for 24 h in Figures 2, 3 and 4, as previously performed [16]. Following 4 days of cell culture, cells were harvested for assays mentioned below. In Figure 1, celastrol (0.1 µM), AZD1981 (10 µM) or DMSO was supplemented freshly every 3 or 4 days over 72 days, and the cumulative population doubling levels (PDLs) were calculated as previously described [7].

**Figure 1.**
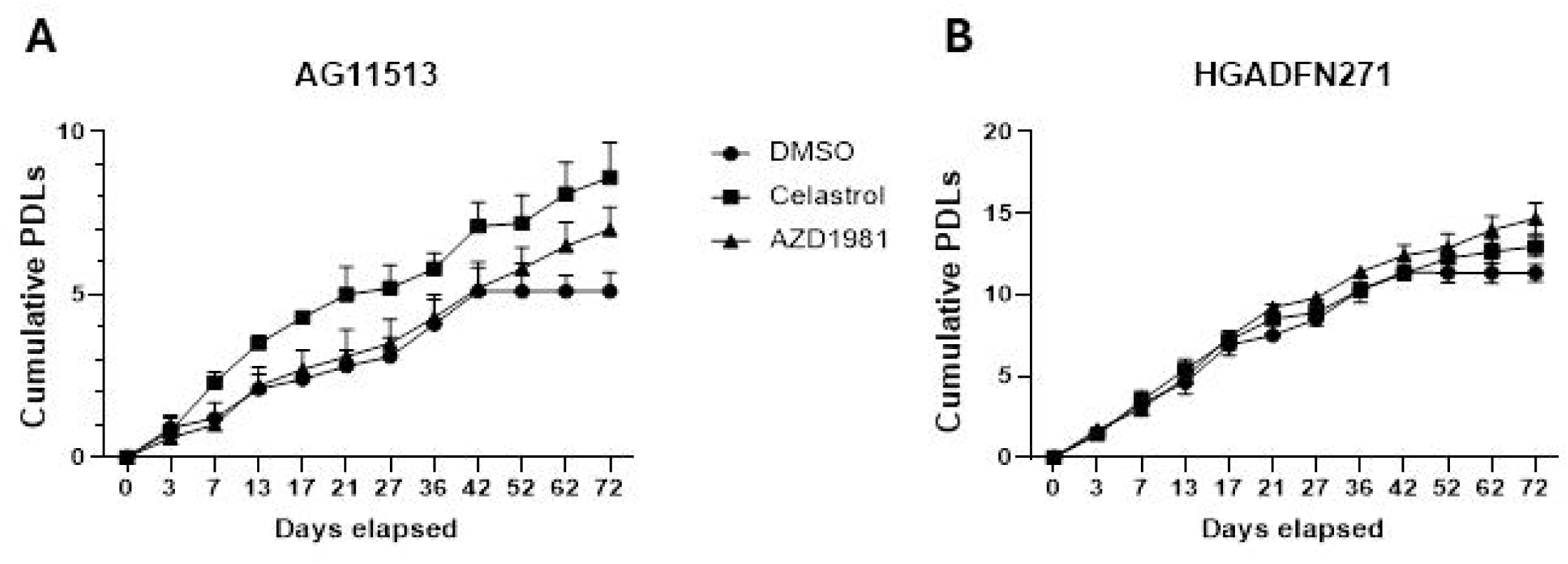
Cell proliferation in HGPS patients-derived fibroblasts AG11513 (A) and HGADFN271 (B) untreated and treated with celastrol or AZD1981. Both fibroblast strains were treated with 0.1 µM celastrol, 10 µM AZD1981, or DMSO alone (untreated control), which was supplemented freshly every 3 or 4 days. Cell proliferation was monitored by periodical cell counting for 72 days and the cumulative population doubling levels (PDLs) were calculated. Note the differences in cumulative PDLs between the two fibroblast strains.

**Figure 2.**
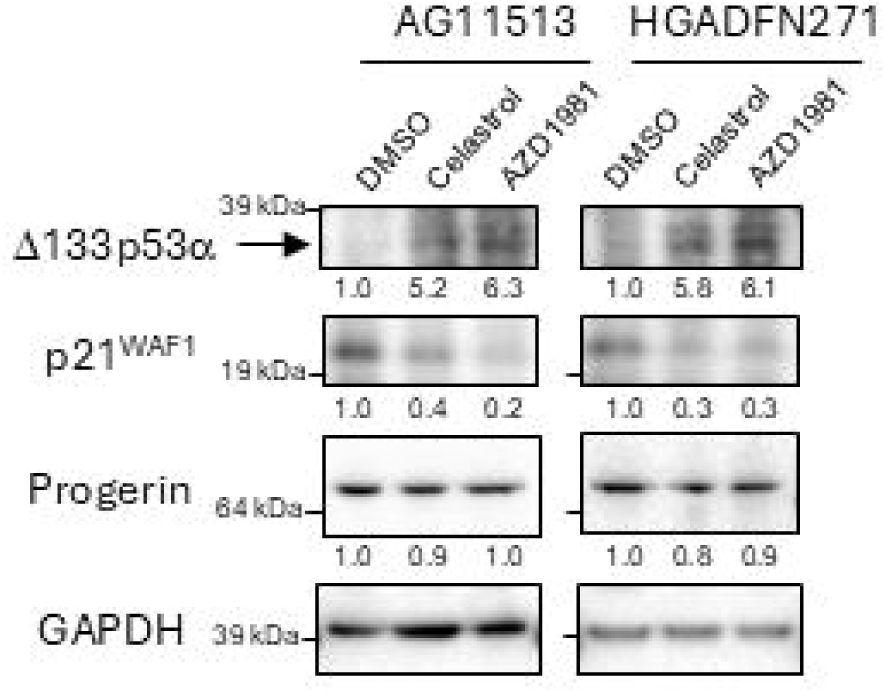
Western blot analysis of HGPS patients-derived fibroblasts AG11513 and HGADFN271 treated with celastrol or AZD1981. The cells were treated with 0.1 µM celastrol, 10 µM AZD1981, or DMSO alone for 24 h, maintained in cell culture for 4 days in the absence of compound, and then harvested. In this figure, Δ133p53α was detected using a rabbit polyclonal antibody MAP4 [2, 30]. The original image obtained with MAP4 before cropping the bands, along with an image obtained using a sheep polyclonal anti-p53 antibody SAPU, which also detects Δ133p53α [1, 30], are shown in Supplementary Figure 1. The quantitative expression levels of Δ133p53α, p21^WAF1^ and progerin, normalized to GAPDH and calculated relative to DMSO alone, are shown below the images.

**Figure 3.**
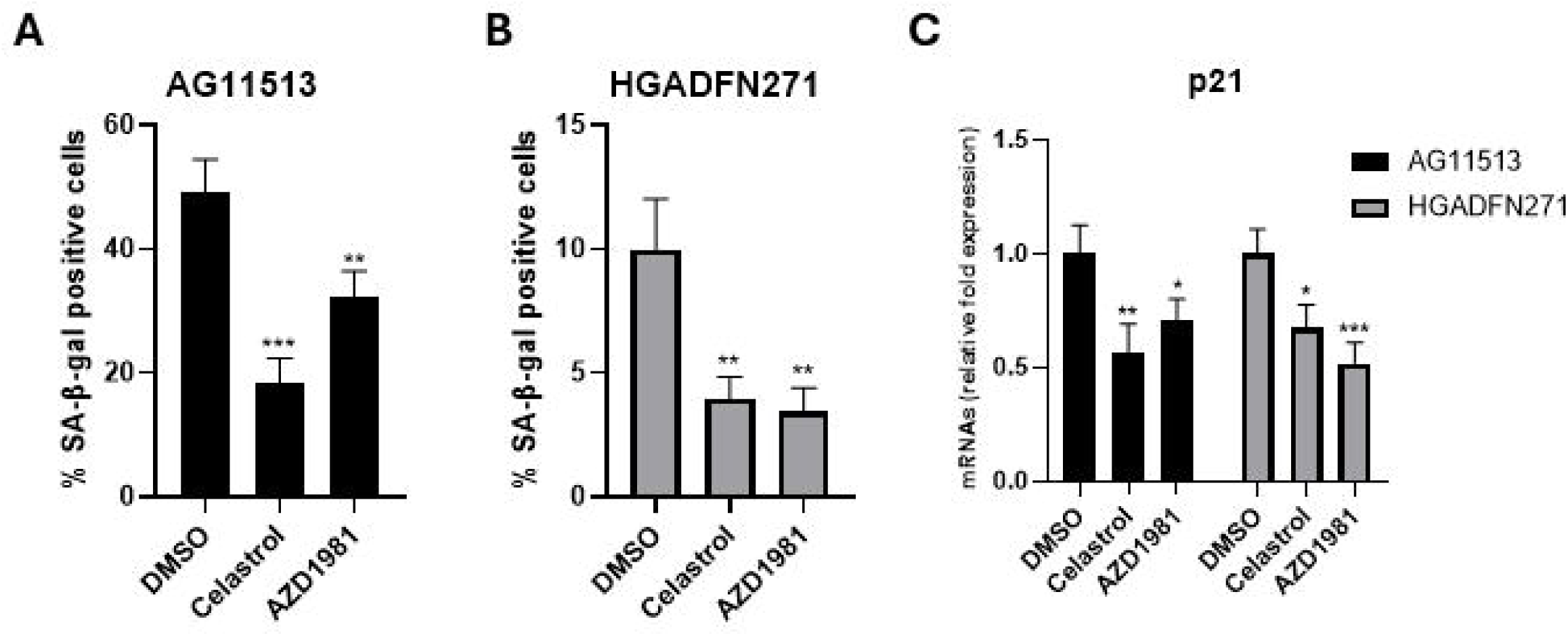
Detection and quantification of senescent cells in HGPS patients-derived fibroblasts treated with celastrol or AZD1981. (**A** and **B**) AG11513 (A) and HGADFN271 (B) fibroblasts were treated as in Figure 2, followed by 4-day cell culture before staining for SA-β-gal activity. The quantitative data for the percentages of SA-β-gal-positive cells are presented as mean ± S.D. obtained from three independent experiments, each with biological triplicates. Note the difference in vertical-axis values between (A) and (B). Representative images are presented in Supplementary Figure 2. (**C**) qRT-PCR assay of p21^WAF1^ mRNA expression. Relative expression levels to DMSO alone are presented as mean ± S.D. from triplicated assays. Unpaired 2-tailed Student’s *t*-test: \**P* < 0.05, \*\**P* < 0.01, \*\*\**P* < 0.001.

**Figure 4.**
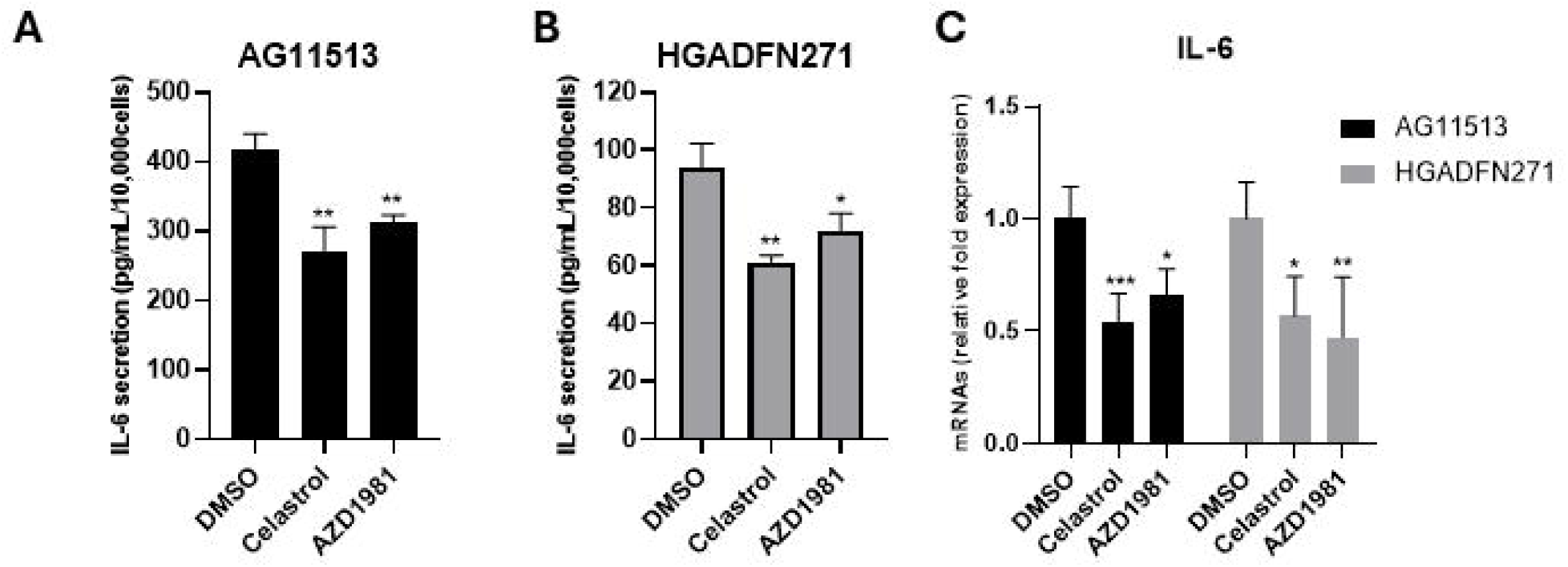
Quantification of IL-6 secretion (A and B) and IL-6 mRNA expression (C) in HGPS patients-derived fibroblasts AG11513 and HGADFN271 treated with celastrol or AZD1981. The cells were treated as in Figure 2. (**A** and **B**) Following 4-day cell culture, culture media were collected for IL-6 ELISA and cell numbers were counted for normalization. The concentrations of secreted IL-6 (pg/ml) were normalized to cell numbers (per 1 × 10^4^) and are presented as mean ± S.D. from biological triplicates. Note the difference in vertical-axis values between AG11513 (A) and HGADFN271 (B). (**C**) qRT-PCR assay of IL-6 mRNA expression. Relative expression levels to DMSO alone are presented as mean ± S.D. from triplicated assays. Unpaired 2-tailed Student’s *t*-test: \**P* < 0.05, \*\**P* < 0.01, \*\*\**P* < 0.001.

### Western blot analysis

Preparation of protein lysates, gel electrophoresis, transfer to PVDF membranes, antibody incubation, and signal detection were performed as previously described [16]. Primary antibodies used were: rabbit polyclonal anti-Δ133p53 antibody MAP4 [2, 30], sheep polyclonal anti-p53 antibody SAPU [1, 30], anti-progerin antibody (Abcam ab153757), anti-p21^WAF1^ antibody (Cell Signaling #2947), and anti-GAPDH antibody (EMD Millipore, MAB374). The quantitative image analysis was performed using the ChemiDoc Imaging System (Bio-Rad) and the Image Lab software (Bio-Rad, ver. 6.1).

### Senescence-associated β-galactosidase (SA-β-gal) staining

SA-β-gal activity was stained using SA-β-Galactosidase Staining Kit (Cell Signaling #9860). Data of SA-β-gal-positive cells (%) were presented as mean ± S.D. from 3 biological replicates in 3 independent experiments. At least 100 cells were observed in each replicate.

### Enzyme-linked immunosorbent assay (ELISA)

Quantification of IL-6 in culture media was carried out using Human IL-6 ELISA Kit (Sigma-Aldrich, RAB0306). Standard curves were drawn from recombinant IL-6 provided in the kit. Data were normalized to cell numbers and presented as mean ± S.D. from biological triplicates.

### RNA isolation and quantitative RT-PCR (qRT-PCR)

Total RNA samples were prepared using RNeasy Plus Micro Kit (QIAGEN #74034). Reverse transcription was carried out using High-Capacity cDNA Reverse Transcription Kit (Thermo Fisher #4368814). qRT-PCR assays were performed using Taqman Gene Expression Master Mix (Thermo Fisher #4369016). The primer/probe sets used were (Thermo Fisher): p21^WAF1^ (Hs00355782_m1), IL-6 (Hs00174131_m1), and GAPDH (Hs02758991_g1) as an internal control. All qRT-PCR data were mean ± S.D. from technical triplicate (n=3).

## RESULTS

### Cell proliferation of two fibroblast strains derived from HGPS patients and the effects of celastrol and AZD1981

In general, primary cells derived from HGPS patients are less proliferative and have a shorter replicative lifespan than those derived from normal donors. However, various factors such as age of patients, methods of strain establishment, and conditions of subsequent cell culture could significantly affect their proliferative potential and replicative lifespan. We therefore monitored two HGPS-derived fibroblast strains, AG11513 and HGADFN271, for long-term proliferation in cell culture (Figure 1A and Figure 1B, DMSO alone). These two strains in our laboratory, at the time of use in this study, were different in proliferative capacity and remaining replicative lifespan: AG11513 fibroblasts (derived from an 8-year-old patient) proliferated slowly and only had ∼5 population doubling levels (PDLs) before proliferation arrest (Figure 1A, DMSO alone), while HGADFN271 fibroblasts (derived from a 1-year-3-month-old patient) still had ∼11 PDLs remaining (Figure 1B, DMSO alone).

Treatment of these fibroblasts with celastrol or AZD1981, added fresh every 3 or 4 days, resulted in continuous proliferation beyond the point at which the control cells became proliferation-arrested (Figure 1A and Figure 1B, after 42 days). This extension of replicative lifespan appeared more pronounced in AG11513 fibroblasts, which had fewer PDLs remaining.

### Celastrol and AZD1981 upregulate endogenous Δ133p53α protein in HGPS-derived fibroblasts

AG11513 and HGADFN271 fibroblasts were treated with celastrol (0.1 µM) or AZD1981 (10 µM) for 24 h, followed by 4 days of cell culture in the absence of compound and then harvested for western blot analysis. Both celastrol and AZD1981 were found to upregulate Δ133p53α protein levels by approximately 5-to 6-fold in these cells (Figure 2 and Supplementary Figure 1), concurrent with decreased expression of p21^WAF1^ (Figure 2), which is known to be repressed by Δ133p53α [2, 7, 15]. Neither compound affected progerin levels (Figure 2).

### Celastrol and AZD1981 reduces cellular senescence in HGPS-derived fibroblasts

Consistent with the proliferative characteristics shown in Figure 1, SA-β-gal-positive senescent cells were more abundant in untreated AG11513 fibroblasts (∼50%, Figure 3A) than in untreated HGADFN271 fibroblasts (∼10%, Figure 3B). Despite this difference in basal senescence levels, treatment with celastrol or AZD1981 significantly reduced the percentage of SA-β-gal-positive senescent cells in both fibroblast strains. In AG11513, celastrol reduced it to ∼20% and AZD1981 to ∼30% (Figure 3A). In HGADFN271, both compounds resulted in only ∼3% of SA-β-gal-positivity (Figure 3B). The mRNA expression levels of p21^WAF1^, a mediator of cellular senescence, were also reduced in these treated fibroblasts (Figure 3C), confirming the protein levels shown in Figure 2. The inhibition of cellular senescence by celastrol and AZD1981 parallels the extension of replicative lifespan by these compounds (Figure 1).

### Celastrol and AZD1981 reduces IL-6 secretion in HGPS-derived fibroblasts

In line with the difference in basal senescence levels (Figure 3A and 3B), the senescence-associated, proinflammatory cytokine IL-6 [31] was secreted into the culture medium at higher levels by untreated AG11513 fibroblasts (Figure 4A) than by untreated HGADFN271 fibroblasts (Figure 4B). Again, despite this difference in basal IL-6 secretion, celastrol and AZD1981 resulted in ∼30% reduction in secreted IL-6 levels in both AG11513 and HGADFN271 fibroblasts (Figure 4A and 4B). The mRNA expression levels of IL-6 were also reduced in these fibroblasts treated with celastrol or AZD1981 (Figure 4C).

## DISCUSSION

This is the first report of testing Δ133p53α-activating compounds in primary human cells from disease patients. Celastrol and AZD1981, when used to treat fibroblasts derived from HGPS patients, were confirmed to upregulate endogenous Δ133p53α levels. These compounds reduced cellular senescence and IL-6 secretion, which are major phenotypes prematurely occurring in these diseased cells, and extended their replicative lifespan. Celastrol and AZD1981 exhibited these effects regardless of the baseline levels of cell proliferation and senescence in these fibroblasts, suggesting that these compounds may act on both actively dividing and proliferation-arrested cells. Consistent with our previous findings with the lentiviral expression of Δ133p53α [2, 7, 15], compound-induced upregulation of Δ133p53α led to repressed expression of p21^WAF1^, in turn leading to inhibition of cellular senescence. It should be noted that celastrol and AZD1981 did not affect the expression level of progerin. Considering that currently available therapies primarily target the synthesis, modification and protein turnover of progerin [32, 33], celastrol and AZD1981, acting independently of progerin regulation, may synergize with these therapies.

Further mechanistic studies are needed for both compounds. While AZD1981 has not been linked to cellular senescence or aging, celastrol has been reported to exert therapeutic effects in mouse and rat models of senescence- and aging-associated diseases and functional decline, such as neurodegenerative diseases [34], osteoarthritis [35], and skeletal muscle atrophy [36]. Since Δ133p53α is present only in humans and primates, but not in mice or rats [5, 37], celastrol could exhibit additional Δ133p53α-dependent therapeutic activities in human diseases beyond those observed in rodent models. A potential synergy between Δ133p53α-activating compounds and senolytic or senomorphic drugs also merits exploration [10].

## CONCLUSION

Celastrol and AZD1981, identified in our previous drug and investigational agent library qHTS, inhibit cellular senescence and IL-6 secretion in fibroblasts derived from HGPS patients, recapitulating the beneficial effects exerted by lentiviral expression of Δ133p53α. These compounds warrant further investigation for potential clinical translation.

## DECLARATIONS

### Authors’ contributions

Performed data acquisition and analyses: Joruiz SM, Lissa D, von Muhlinen N; Made substantial contributions to conception and design of the study: Dranchak PK, Inglese J, Horikawa I, Harris CC; Wrote the manuscript: Joruiz SM, Lissa D, Horikawa I.

### Availability of data and materials

All materials in this study and raw data related to this study are available upon request.

### Financial support and sponsorship

This research was supported by the Intramural Research Program of the National Institutes of Health (NIH): ZIA BC 011496 (Harris CC) and ZIA TR 000052 (Inglese J). The contributions of the NIH authors were made as part of their official duties as NIH federal employees, are in compliance with agency policy requirements, and are considered Works of the United States Government. However, the findings and conclusions presented in this paper are those of the authors and do not necessarily reflect the views of the NIH or the U.S. Department of Health and Human Services.

### Conflicts of interest

Lissa D is a current employee at AstraZeneca but had no association with the company at the time of this project, and no participation from AstraZeneca was received in any form. All authors declared that there are no conflicts of interest.

### Ethical approval and informed consent

Not applicable.

## Notes

### Competing Interest Statement

The authors have declared no competing interest.

### Summary of Updates

Figure 2 revised with additional image quantification. Supplementary Figure 2 added.

## References

1. Joruiz SM, Bourdon JC. p53 isoforms: Key regulators of the cell fate decision. Cold Spring Harb Perspect Med, 2016, 6(8): a026039. doi: 10.1101/cshperspect.a026039

2. Fujita K, Mondal AM, Horikawa I, Nguyen GH, Kumamoto K, Sohn JJ, et al. p53 isoforms Δ133p53 and p53β are endogenous regulators of replicative cellular senescence. Nat Cell Biol, 2009, 11(9): 1135–1142. doi: 10.1038/ncb1928

3. Mondal AM, Horikawa I, Pine SR, Fujita K, Morgan KM, Vera E, et al. p53 isoforms regulate aging- and tumor-associated replicative senescence in T lymphocytes. J Clin Invest, 2013, 123(12): 5247–5257. doi: 10.1172/jci70355

4. Horikawa I, Fujita K, Jenkins LM, Hiyoshi Y, Mondal AM, Vojtesek B, et al. Autophagic degradation of the inhibitory p53 isoform Δ133p53α as a regulatory mechanism for p53-mediated senescence. Nat Commun, 2014, 5: 4706. doi: 10.1038/ncomms5706

5. Turnquist C, Horikawa I, Foran E, Major EO, Vojtesek B, Lane DP, et al. p53 isoforms regulate astrocyte-mediated neuroprotection and neurodegeneration. Cell Death Differ, 2016, 23(9): 1515–1528. doi: 10.1038/cdd.2016.37

6. Mondal AM, Zhou H, Horikawa I, Suprynowicz FA, Li G, Dakic A, et al. Δ133p53α, a natural p53 isoform, contributes to conditional reprogramming and long-term proliferation of primary epithelial cells. Cell Death Dis, 2018, 9(7): 750. doi: 10.1038/s41419-018-0767-7

7. von Muhlinen N, Horikawa I, Alam F, Isogaya K, Lissa D, Vojtesek B, et al. p53 isoforms regulate premature aging in human cells. Oncogene, 2018, 37(18): 2379–2393. doi: 10.1038/s41388-017-0101-3

8. Turnquist C, Beck JA, Horikawa I, Obiorah IE, von Muhlinen N, Vojtesek B, et al. Radiation-induced astrocyte senescence is rescued by Δ133p53. Neuro Oncol, 2019, 21(4): 474–485. doi: 10.1093/neuonc/noz001

9. Ungerleider K, Beck JA, Lissa D, Joruiz S, Horikawa I, Harris CC. Δ133p53α protects human astrocytes from amyloid-beta induced senescence and neurotoxicity. Neuroscience, 2022, 498: 190–202. doi: 10.1016/j.neuroscience.2022.06.004

10. Horikawa I, Yamada L, Harris BT, Harris CC. Δ133p53α-mediated inhibition of astrocyte senescence and neurotoxicity as a possible therapeutic approach for neurodegenerative diseases. Neuroscience, 2025, 580: 54–61.

11. Roselle C, Horikawa I, Chen L, Kelly AR, Gonzales D, Da T, et al. Enhancing chimeric antigen receptor T cell therapy by modulating the p53 signaling network with Δ133p53α. Proc Natl Acad Sci U S A, 2024, 121(10): e2317735121. doi: 10.1073/pnas.2317735121

12. Donehower LA, Harvey M, Slagle BL, McArthur MJ, Montgomery CA, Jr., Butel JS, et al. Mice deficient for p53 are developmentally normal but susceptible to spontaneous tumours. Nature, 1992, 356(6366): 215–221. doi: 10.1038/356215a0

13. Efeyan A, Serrano M. p53: guardian of the genome and policeman of the oncogenes. Cell Cycle, 2007, 6(9): 1006–1010. doi: 10.4161/cc.6.9.4211

14. Horikawa I, Harris CC. Δ133p53: A p53 isoform enriched in human pluripotent stem cells. Cell Cycle, 2017, 16: 1631–1632. doi: 10.1080/15384101.2017.1345228

15. Horikawa I, Park KY, Isogaya K, Hiyoshi Y, Li H, Anami K, et al. Δ133p53 represses p53-inducible senescence genes and enhances the generation of human induced pluripotent stem cells. Cell Death Differ, 2017, 24(6): 1017–1028. doi: 10.1038/cdd.2017.48

16. Lissa D, Joruiz SM, Dranchak PK, Ungerleider K, Yamada L, Horikawa I, et al. A quantitative high-throughput screen identifies compounds that upregulate the p53 isoform Δ133p53α and inhibit cellular senescence. ACS Pharmacol Transl Sci, 2025, 10.1021/acsptsci.5c00186.

17. Salminen A, Lehtonen M, Paimela T, Kaarniranta K. Celastrol: Molecular targets of Thunder God Vine. Biochem Biophys Res Commun, 2010, 394(3): 439–442. doi: 10.1016/j.bbrc.2010.03.050

18. Seo HR, Seo WD, Pyun BJ, Lee BW, Jin YB, Park KH, et al. Radiosensitization by celastrol is mediated by modification of antioxidant thiol molecules. Chem Biol Interact, 2011, 193(1): 34–42. doi: 10.1016/j.cbi.2011.04.009

19. Trott A, West JD, Klaić L, Westerheide SD, Silverman RB, Morimoto RI, et al. Activation of heat shock and antioxidant responses by the natural product celastrol: transcriptional signatures of a thiol-targeted molecule. Mol Biol Cell, 2008, 19(3): 1104–1112. doi: 10.1091/mbc.e07-10-1004

20. Westerheide SD, Bosman JD, Mbadugha BN, Kawahara TL, Matsumoto G, Kim S, et al. Celastrols as inducers of the heat shock response and cytoprotection. J Biol Chem, 2004, 279(53): 56053–56060. doi: 10.1074/jbc.M409267200

21. Yang H, Chen D, Cui QC, Yuan X, Dou QP. Celastrol, a triterpene extracted from the Chinese “Thunder of God Vine,” is a potent proteasome inhibitor and suppresses human prostate cancer growth in nude mice. Cancer Res, 2006, 66(9): 4758–4765. doi: 10.1158/0008-5472.Can-05-4529

22. Schmidt JA, Bell FM, Akam E, Marshall C, Dainty IA, Heinemann A, et al. Biochemical and pharmacological characterization of AZD1981, an orally available selective DP2 antagonist in clinical development for asthma. Br J Pharmacol, 2013, 168(7): 1626–1638. doi: 10.1111/bph.12053

23. Eriksson M, Brown WT, Gordon LB, Glynn MW, Singer J, Scott L, et al. Recurrent de novo point mutations in lamin A cause Hutchinson-Gilford progeria syndrome. Nature, 2003, 423(6937): 293–298. doi: 10.1038/nature01629

24. Hennekam RC. Hutchinson-Gilford progeria syndrome: review of the phenotype. Am J Med Genet A, 2006, 140(23): 2603–2624. doi: 10.1002/ajmg.a.31346

25. Ding SL, Shen CY. Model of human aging: recent findings on Werner’s and Hutchinson-Gilford progeria syndromes. Clin Interv Aging, 2008, 3(3): 431–444.

26. Brassard JA, Fekete N, Garnier A, Hoesli CA. Hutchinson-Gilford progeria syndrome as a model for vascular aging. Biogerontology, 2016, 17(1): 129–145. doi: 10.1007/s10522-015-9602-z

27. Wheaton K, Campuzano D, Ma W, Sheinis M, Ho B, Brown GW, et al. Progerin-Induced Replication Stress Facilitates Premature Senescence in Hutchinson-Gilford Progeria Syndrome. Mol Cell Biol, 2017, 37(14): e00659. doi: 10.1128/mcb.00659-16

28. Xu Q, Mojiri A, Boulahouache L, Morales E, Walther BK, Cooke JP. Vascular senescence in progeria: role of endothelial dysfunction. Eur Heart J Open, 2022, 2(4): oeac047. doi: 10.1093/ehjopen/oeac047

29. Batista NJ, Desai SG, Perez AM, Finkelstein A, Radigan R, Singh M, et al. The Molecular and Cellular Basis of Hutchinson-Gilford Progeria Syndrome and Potential Treatments. Genes (Basel), 2023, 14(3): 602. doi: 10.3390/genes14030602

30. Marcel V, Khoury MP, Fernandes K, Diot A, Lane DP, Bourdon JC. Detecting p53 isoforms at protein level. Methods Mol Biol, 2013, 962: 15–29. doi: 10.1007/978-1-62703-236-0_2

31. Ungerleider K, Beck J, Lissa D, Turnquist C, Horikawa I, Harris BT, et al. Astrocyte senescence and SASP in neurodegeneration: Tau joins the loop. Cell Cycle, 2021, 20(8): 752–764. doi: 10.1080/15384101.2021.1909260

32. Arun A, Nath AR, Thankachan B, Unnikrishnan MK. Hutchinson-Gilford progeria syndrome: unraveling the genetic basis, symptoms, and advancements in therapeutic approaches. Ther Adv Rare Dis, 2024, 5: 26330040241305144. doi: 10.1177/26330040241305144

33. Koblan LW, Erdos MR, Wilson C, Cabral WA, Levy JM, Xiong ZM, et al. In vivo base editing rescues Hutchinson-Gilford progeria syndrome in mice. Nature, 2021, 589(7843): 608–614. doi: 10.1038/s41586-020-03086-7

34. Liu D, Zhang Q, Luo P, Gu L, Shen S, Tang H, et al. Neuroprotective Effects of Celastrol in Neurodegenerative Diseases-Unscramble Its Major Mechanisms of Action and Targets. Aging Dis, 2022, 13(3): 815–836. doi: 10.14336/ad.2021.1115

35. Yang G, Wang K, Song H, Zhu R, Ding S, Yang H, et al. Celastrol ameliorates osteoarthritis via regulating TLR2/NF-κB signaling pathway. Front Pharmacol, 2022, 13: 963506. doi: 10.3389/fphar.2022.963506

36. Yadav A, Yadav SS, Singh S, Dabur R. Natural products: Potential therapeutic agents to prevent skeletal muscle atrophy. Eur J Pharmacol, 2022, 925: 174995. doi: 10.1016/j.ejphar.2022.174995

37. Joruiz SM, Beck JA, Horikawa I, Harris CC. The Δ133p53 Isoforms, tuners of the p53 pathway. Cancers (Basel), 2020, 12(11): 3422. doi: 10.3390/cancers12113422

