## Supplementary Figures 1 and 2 for "Pharmacologic activation of Δ133p53α reduces cellular senescence in progeria patients-derived cells"

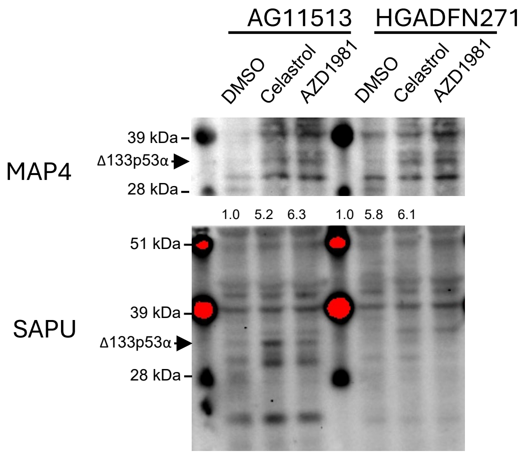


**Supplementary Figure 1. Original western blot images obtained with MAP4 and SAPU antibodies.** The images obtained with the rabbit polyclonal antibody MAP4 (above; cropped bands shown in Figure 2) and the sheep polyclonal antibody SAPU (below) both consistently showed bands corresponding to Δ133p53α, which were upregulated by celastrol and AZD1981 (indicated by arrows).

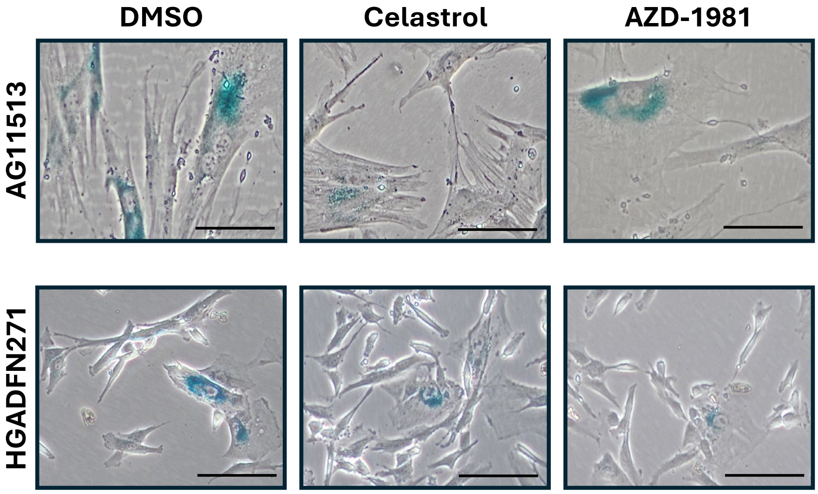


**Supplementary Figure 2. Representative images of SA-β-gal staining.** Magnification, ×40. Scale bars, 50 μm.
